# Estimating multimodal brain variability in schizophrenia spectrum disorders: A worldwide ENIGMA study

**DOI:** 10.1101/2023.09.22.559032

**Authors:** Wolfgang Omlor, Finn Rabe, Simon Fuchs, Giacomo Cecere, Stephanie Homan, Werner Surbeck, Nils Kallen, Foivos Georgiadis, Tobias Spiller, Erich Seifritz, Thomas Weickert, Jason Bruggemann, Cynthia Weickert, Steven Potkin, Ryota Hashimoto, Kang Sim, Kelly Rootes-Murdy, Yann Quide, Josselin Houenou, Nerisa Banaj, Daniela Vecchio, Fabrizio Piras, Federica Piras, Gianfranco Spalletta, Raymond Salvador, Andriana Karuk, Edith Pomarol-Clotet, Amanda Rodrigue, Godfrey Pearlson, David Glahn, David Tomecek, Filip Spaniel, Antonin Skoch, Matthias Kirschner, Stefan Kaiser, Peter Kochunov, Feng-Mei Fan, Ole A. Andreassen, Lars T. Westlye, Pierre Berthet, Vince D Calhoun, Fleur Howells, Anne Uhlmann, Freda Scheffler, Dan Stein, Felice Iasevoli, Murray J. Cairns, Vaughan J. Carr, Stanley V. Catts, Maria A. Di Biase, Assen Jablensky, Melissa J. Green, Frans A. Henskens, Paul Klauser, Carmel Loughland, Patricia T. Michie, Bryan Mowry, Christos Pantelis, Paul E. Rasser, Ulrich Schall, Rodney Scott, Andrew Zalesky, Andrea de Bartolomeis, Annarita Barone, Mariateresa Ciccarelli, Arturo Brunetti, Sirio Cocozza, Giuseppe Pontillo, Mario Tranfa, Annabella Di Giorgio, Sophia I. Thomopoulos, Neda Jahanshad, Paul M. Thompson, Theo van Erp, Jessica Turner, Philipp Homan

**Affiliations:** Psychiatric Hospital, University of Zurich, Zurich, Switzerland; Neuroscience Center Zurich, University of Zurich & Swiss Federal Institute of Technology Zurich, Zurich, Switzerland; Department of Neuroscience and Physiology, Upstate Medical University, Syracuse, New York, United States; Edith Collins Centre, Sydney Local Health District, Sydney 2050, Australia; Clinical Translational Neuroscience Laboratory, Department of Psychiatry and Human Behavior, University of California Irvine, Irvine, CA 92697; Department of Pathology of Mental Diseases, National Center of Neurology and Psychiatry, National Institute of Mental Health Tokyo 187-8551, Japan; Institute of Mental Health, Singapore 539747, Singapore; Psychology Department and Neuroscience Institute, Georgia State University, Atlanta, GA 30303; School of Psychology, UNSW Sydney, Sydney, Australia; Hopitaux Universitaires Henri Mondor, Crétail, France; Laboratory of Neuropsychiatry, Clinical Neuroscience and Neurorehabilitation Department, Santa Lucia Foundation IRCCS, Rome, Italy; FIDMAG Sisters Hospitaliers Research Foundation, Barcelona, Spain; Department of Psychiatry, Boston’s Children Hospital Harvard Medical School, Boston, MA 02115; Department of Psychiatry, Yale University, New Haven, Connecticut; Olin Neuropsychiatric Research Center, Institute of Living, Hartford Hospital, Hartford, Connecticut; National Institute of Mental Health, Klecany, Czech Republic; Division of Adult Psychiatry, Department of Psychiatry, Geneva University Hospitals, Geneva, Switzerland; Maryland Psychiatric Research Center Department of Psychiatry, University of Maryland, School of Medicine, Baltimore, MD, USA; Beijing Huilongguan Hospital Peking University Huilongguan Clinical Medical School, Beijing, China; NORMENT Centre, Division of Mental Health and Addiction, University of Oslo and Oslo University Hospital, Oslo, Norway, and KG JEbsen Centre for Neurodevelopment, University of Oslo, Oslo, Norway; Neuroscience Institute University of Cape Town, Cape Town, South Africa; Department of Psychiatry and Mental Health University of Cape Town, Cape Town, South Africa; Department of Child and Adolescent Psychiatry Technische Universität Dresden, Dresden 01187, Germany; SA MRC Unit on Risk and Resilience in Mental Disorders Department of Psychiatry, Neuroscience Insitute University of Cape Town, Cape Town, South Africa; Laboratory of Molecular and Translational Psychiatry and Unit of Treatment Resistant Psychosis, Section of Psychiatry, Reproductive and Odontostomatological Sciences, Department of Neuroscience, University of Naples Federico II, Naples, Italy; Neuroscience Research Australia, Sydney 2031, Australia; School of Psychiatry, University of New South Wales, Sydney 2033, Australia; Specialty of Addiction Medicine, Central Clinical School, Faculty of Medicine and Health, University of Sydney, Sydney 2006, Australia; Center for the Neurobiology of Learning and Memory, University of California Irvine, Irvine, CA 92697; Department of Psychiatry and Behavioral Health, Wexner Medical Center, The Ohio State University, Columbus, OH, USA; Tri-institutional Center for Translational Research in Neuroimaging and Data Science (TReNDS), Georgia State University, Georgia Institute of Technology, Emory University, Atlanta, GA, USA; School of Biomedical Sciences and Pharmacy, University of Newcastle, Callaghan, NSW, Australia; Centre for Brain and Mental Health Research, University of Newcastle, Callaghan, NSW, Australia; Hunter Medical Research Institute, New Lambton Heights, NSW, Australia; School of Clinical Medicine, Discipline of Psychiatry and Mental Health, The University of New South Wales (UNSW), Sydney, NSW, Australia; Neuroscience Research Australia, Randwick, NSW, Australia; Department of Psychiatry, Monash University, Clayton, VIC, Australia; School of Medicine, University of Queensland, Brisbane, Australia; Melbourne Neuropsychiatry Centre, Department of Psychiatry, the University of Melbourne & Melbourne Health, Carlton South, VIC, Australia; Department of Psychiatry, Brigham and Women’s Hospital, Harvard Medical School, Boston, MA, USA; University of Western Australia, Perth, WA, Australia; School of Medicine & Public Health, The University of Newcastle, Newcastle, NSW, Australia; Priority Research Centre for Health Behaviour, the University of Newcastle, Newcastle, NSW, Australia; Hunter Medical Research Institute, Newcastle, NSW, Australia; Brain and Mental Health Laboratory, Monash Institute of Cognitive and Clinical Neurosciences, School of Psychological Sciences and Monash Biomedical Imaging, Monash University, Clayton, VIC, Australia; Department of Psychiatry, Lausanne University Hospital and the University of Lausanne, Lausanne, Switzerland; School of Psychological Sciences, University of Newcastle, Callaghan, NSW, Australia; Queensland Brain Institute, the University of Queensland, Brisbane, QLD, Australia; The Queensland Centre for Mental Health Research, the University of Queensland, Brisbane, QLD, Australia; Florey Institute of Neuroscience & Mental Health, Parkville, VIC, Australia; School of Medicine and Public Health, College of Health, Medicine and Wellbeing, The University of Newcastle, Callaghan, NSW, Australia; Melbourne School of Engineering, the University of Melbourne, Parkville, VIC, Australia; Section of Psychiatry - Department of Neuroscience, University Federico II, Naples, Italy; Department of Advanced Biomedical Sciences, University Federico II, Naples, Italy; Department of Electrical Engineering and Information Technology (DIETI), University Federico II, Naples, Italy; ASST Papa Giovanni XXIII, Mental Health Department, Bergamo, Italy; Imaging Genetics Center, Stevens Neuroimaging & Informatics Institute, Keck School of Medicine, University of Southern California, Los Angeles, CA, USA

**Author notes:** Correspondence concerning this article should be addressed to Philipp Homan, MD, PhD, Psychiatric Hospital, University of Zurich, Lenggstrasse 31, 8032 Zurich, Switzerland.

## Abstract

**Objective:** Schizophrenia is a multifaceted disorder associated with structural brain heterogeneity. Despite its relevance for identifying illness subtypes and informative biomarkers, structural brain heterogeneity in schizophrenia remains incompletely understood. Therefore, the objective of this study was to provide a comprehensive insight into the structural brain heterogeneity associated with schizophrenia.

**Methods:** This meta- and mega-analysis investigated the variability of multimodal structural brain measures of white and gray matter in individuals with schizophrenia versus healthy controls. Using the ENIGMA dataset of MRI-based brain measures from 22 international sites with up to 6139 individuals for a given brain measure, we examined variability in cortical thickness, surface area, folding index, subcortical volume and fractional anisotropy.

**Results:** We found that individuals with schizophrenia are distinguished by higher heterogeneity in the frontotemporal network with regard to multimodal structural measures. Moreover, individuals with schizophrenia showed higher homogeneity of the folding index, especially in the left parahippocampal region.

**Conclusions:** Higher multimodal heterogeneity in frontotemporal regions potentially implies different subtypes of schizophrenia that converge on impaired frontotemporal interaction as a core feature of the disorder. Conversely, more homogeneous folding patterns in the left parahippocampal region might signify a consistent characteristic of schizophrenia shared across subtypes. These findings underscore the importance of structural brain variability in advancing our neurobiological understanding of schizophrenia, and aid in identifying illness subtypes as well as informative biomarkers.

## Introduction

Schizophrenia is characterized by an array of biological and symptomatic heterogeneity that remains incompletely understood^1–3^. Notably, the biological heterogeneity of schizophrenia is reflected in structural irregularities of the brain^4–8^ as well as functional abnormalities^9,10^. Historically, neuroimaging studies including meta-analyses have predominantly focused on evaluating mean differences in structural brain measures between individuals with psychosis and healthy controls^11–13^. However, recent studies also emphasized that individuals with or at clinical high risk for psychosis exhibit differences in the variability of structural brain measures compared to healthy controls^7,14–18^: While higher variability of specific brain structures may be associated with schizophrenia subtypes, lower variability argues for a more stable feature of the disorder that is shared across illness subtypes^7^.

Therefore, a shift in perspective towards the analysis of variability beyond mean differences is important to guide neurobiological research on schizophrenia, with clinical relevance for identifying illness subtypes and biomarkers as well as for analyzing medication effects^19–21^. In addition to regional structural measures, global measures such as the person-based similarity index (PBSI)^15,22^ have been applied, for example, to reveal higher divergence of general cortical thickness, surface area and subcortical volume in individuals at high risk for psychosis^17^. Against this backdrop, our meta-analysis study investigated a variety of MRI-based brain measures, both for gray and white matter structures and at both global and regional levels, to provide a comprehensive, multimodal assessment of neuroanatomical heterogeneity in schizophrenia. To overcome limitations of individual studies with small sample sizes and to support the replicability of our findings, multimodal variability was explored using meta- and mega-analysis of a large dataset from the Enhancing Neuroimaging Genetics through Meta-Analysis (ENIGMA) consortium^13,23^.

## Methods

### Participants

The ENIGMA dataset comprised neuroimaging data from 22 international sites, including 15 sites with individual-level data and 7 sites with group-level data. Sites with individual-level data provided cortical, subcortical and white matter measures for individuals with schizophrenia and healthy controls, while sites with group-level data provided means and standard deviations of respective measures across individuals with schizophrenia and healthy controls. The dataset included 6138 individuals for cortical thickness, 6138 for cortical surface area, 5701 for cortical folding index, 6139 for subcortical volume and 2636 individuals for white matter measure fractional anisotropy. Prior to data collection, each site obtained ethics committee approval, and participants provided informed consent or assent prior to participation.

### MRI processing

Neuroimaging data were processed according to the standardized ENIGMA protocols (http://enigma.ini.usc.edu/protocols/imaging-protocols/). Motion correction (DTI analysis), automated Talairach transformation, skull stripping, segmentation of the subcortical white and gray matter volumetric structures as well as intensity normalisation were incorporated by automated FreeSurfer pipelines^24–31^. Preprocessing procedures generated values of fractional anisotropy for white matter labels, values of subcortical volumes as well as values of cortical thickness, surface area and folding index for cortical regions according to the Desikan-Killiany atlas^32^. As part of the ENIGMA quality control procedure, outliers (±2 SD from the mean) were identified and all images were visually inspected to exclude poorly segmented regions from the analyses. These quality control procedures led to minor differences in sample sizes for each region of interest (ROI). The application of this protocol finally yielded 68 cortical ROIs, 18 subcortical ROIs and 63 DTI ROIs. Subjects younger than 18 years or older than 65 years were excluded from further analyses.

### Statistical analysis

#### Variability ratio

Statistical analyses were performed in R version 4.2.0 and Python version 3.9.16. Variability ratio (VR) was computed and visualized as forest plots in R using the packages *metafor* and *tidyverse*. To compare variability in regional neuroanatomical measures between individuals with schizophrenia and healthy controls, the natural logarithm of VR was calculated for each ROI according to the following formula:

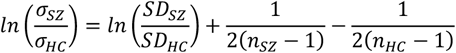

where σ is the unbiased estimate of the standard deviation, SD the reported sample standard deviation and *n* the sample size for the respective group. The results of the VR analysis were summarized in forest plots which exhibit the VR with 95% confidence intervals across all ROIs for a given measure (cortical thickness, surface area, folding index, fractional anisotropy and subcortical volume). For the forest plots, log-transformed VR values were back-transformed onto a linear scale (VR) to facilitate interpretation: VR of 1 indicates equal variability in neuroanatomical measures between schizophrenia patients and healthy controls, while VR > 1 indicates greater and VR < 1 lower variability in schizophrenia patients compared to controls, respectively. Brain surface plots in which the VR for cortical thickness, surface area, folding index, fractional anisotropy and subcortical volume is illustrated with color-coded z-values were generated in Python, using the libraries *mayavi, nibabel, surfer* and *matplotlib*.

#### Coefficient of variation ratio

In biological systems, variance usually scales with the mean such that larger mean values are coupled with larger variance^33^. For a given structure and measure, a between-group difference in variability could therefore be driven by a between-group difference in mean values. For each measure, we therefore also investigated how mean and standard deviation (SD) differences between patients and controls varied across sites. While most of the cortical, subcortical and white matter regions showed correlations between mean and SD differences close to zero, some regions showed moderate positive correlations (**Supplementary Fig. 1**). Even though there were also a few regions with negative correlations between mean and SD differences, the magnitudes of these negative correlations was comparably low (**Supplementary Fig. 2**). To assess the potential impact of correlations between mean and SD on our variability ratio results, we also computed the coefficient of variation ratio (CVR) which incorporates between-group differences in mean to quantify between-group differences in variability^7^:

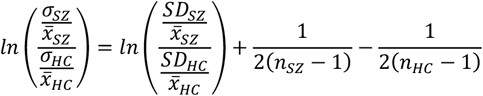

where 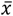 is the mean for the respective group. As for VR, the CVR analysis was illustrated using forest plots. For the forest plots, log-transformed CVR values were back-transformed onto a linear scale and brain surface plots with color-coded CVR z-values were generated.

#### Person-based similarity index

In addition to the region-specific VR and CVR analyses, we calculated the person-based similarity index (PBSI) to explore between-group differences of variability for a given measure across brain regions^16,22^. Separately for the brain measures cortical thickness, surface area, folding index, fractional anisotropy and subcortical volume, the PBSI was computed for the patient and control group according to the formula:

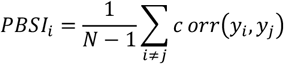

For *j* ≠ *i*, the PBSI of the *i*th participant is the mean Spearman correlation between their brain measure *y*_*i*_ and the brain measure *y*_*j*_ of any other participant of the same group with *N* individuals. For a given participant *i, y*_*i*_ represents a vector in which all of their values for a given measure (e.g. cortical thickness) are concatenated. With regard to a given brain measure, each participant’s PBSI score therefore indicates their similarity to all other members of the same group (PBSI scores closer to 1 indicate higher similarity). The PBSI scores for cortical thickness, surface area, folding index, subcortical volume and fractional anisotropy were then compared between individuals with schizophrenia and healthy controls using a Mann-Whitney *U* test with Bonferroni correction.

## Results

### Cortical thickness

Compared to controls, the schizophrenia group exhibited greater variability in bilateral inferior, middle and superior temporal regions as well as in bilateral superiorfrontal regions. From these frontotemporal regions, the largest effect sizes were found in the right middle (VR = 1.19, 95% CI: 1.08 - 1.30) and left superior temporal areas (VR = 1.16, 95% CI: 1.10 - 1.22). Unilaterally, the schizophrenia group demonstrated greater variability in the left supramarginal region (VR = 1.11, 95% CI: 1.03 - 1.19), in the right pars orbitalis region (VR = 1.07, 95% CI: 1.01 - 1.13), in the left fusiform region (VR = 1.09, 95% CI: 1.01 - 1.17) and in the right precentral region (VR = 1.10, 95% CI: 1.01 - 1.21). Compared to individuals with schizophrenia, healthy controls did not show significantly higher variability in cortical thickness for any cortical region (**Fig. 1**). When the coefficient of variation ratio was computed instead of the variability ratio, all regions with higher heterogeneity in schizophrenia remained significantly different from controls, and controls still did not show higher heterogeneity in any region (**Supplementary Fig. 3**).

**Figure 1:**
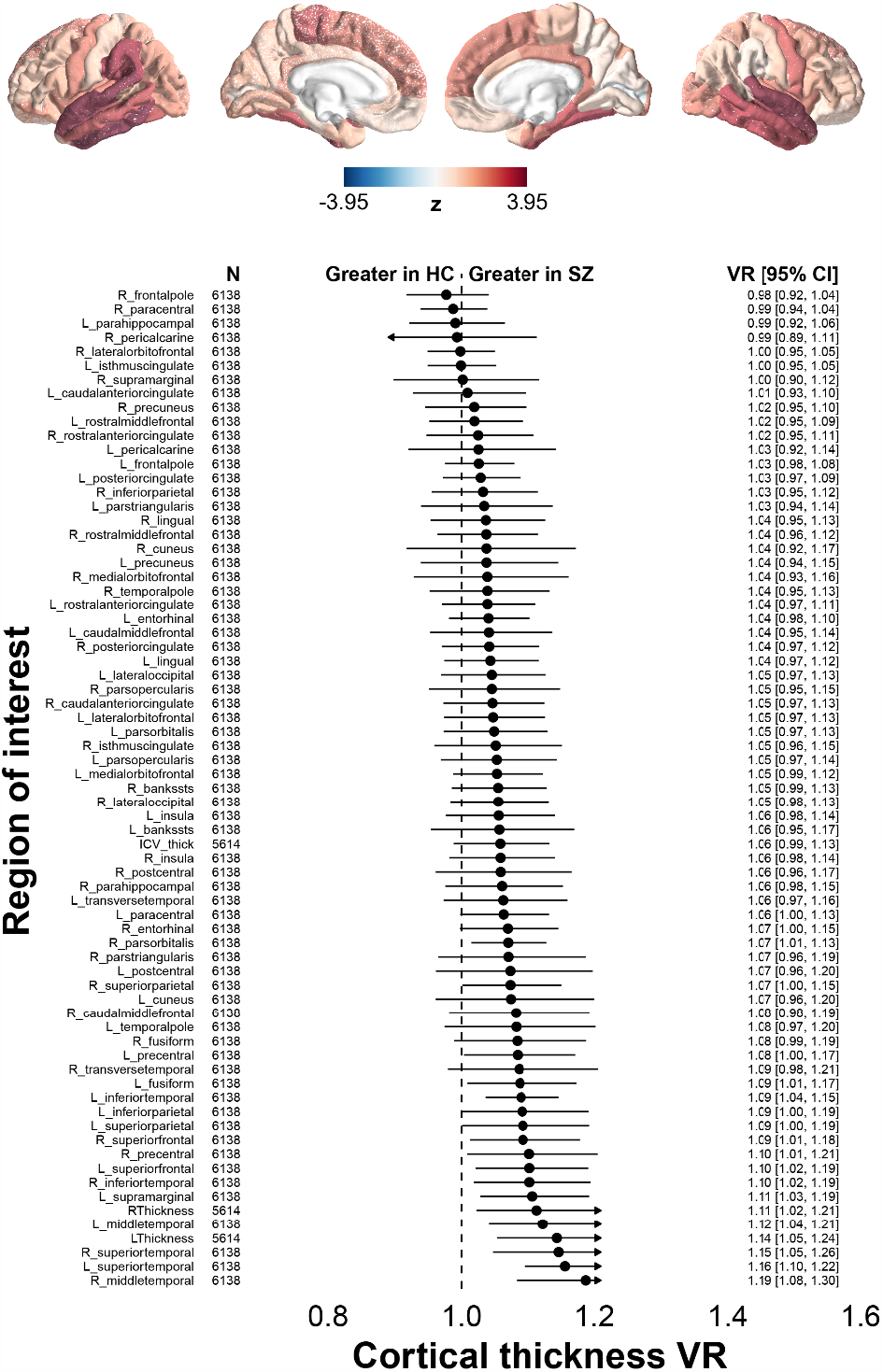
Variability ratio of cortical thickness. Lower panel: Variability ratio (VR) effect sizes for individuals with schizophrenia (SZ) vs. healthy controls (HC) are shown for different cortical regions and on a linear scale, statistically controlling for age and gender. Upper panel: VR effect sizes are projected onto the brain surface and color-coded as z-values. In these cortical maps, areas with higher VRs in patients appear in gradients of red, and areas with higher VRs in controls appear in gradients of blue. CI: Confidence Interval. Independent of the applied variability measure (variability ratio vs. coefficient of variation ratio), higher heterogeneity in schizophrenia was observed for the following regions: Bilateral superior, middle and inferior temporal region, bilateral superior frontal region, left supramarginal and fusiform region as well as right pars orbitalis and precentral region.

### Cortical surface area

Compared with controls, greater variability in the schizophrenia group was particularly evident in the following regions: Right superiortemporal region (VR = 1.12, 95% CI: 1.02 - 1.23), cortical areas around the right superior temporal sulcus (VR = 1.11, 95% CI: 1.03 - 1.20) and right supramarginal region (VR = 1.11, 95% CI: 1.04 - 1.18, **Fig. 2**). Moreover, greater variability in the schizophrenia group was found in the left postcentral region (VR = 1.09, 95% CI: 1.01 - 1.17), in the right medial orbitofrontal region (VR = 1.09, 95% CI: 1.02 - 1.15), in the left superiorparietal region (VR = 1.07, 95% CI: 1.01 - 1.14), in the right middle temporal region (VR = 1.09, 95% CI: 1.01 - 1.17) as well as in the left precuneus (VR = 1.06, 95% CI: 1.01 - 1.11) and right lingual region (VR = 1.10, 95% CI: 1.01 - 1.19, **Fig. 2**). Bilaterally, the superior frontal and transverse temporal areas showed higher variability in comparison to healthy controls (left superior frontal region: VR = 1.07, 95% CI: 1.01 - 1.12; right superior frontal region: VR = 1.06, 95% CI: 1.01 - 1.12; left transverse temporal region: VR = 1.07, 95% CI: 1.02 - 1.13; right transverse temporal region: VR = 1.08, 95% CI: 1.01 - 1.16, **Fig. 2**). When compared to the schizophrenia group, the control group did not exhibit higher variability of the cortical surface area in any cortical region (**Fig. 2**). When we computed the coefficient of variation instead of the variability ratio, all regions with higher variability in schizophrenia remained significantly different from controls, and again no regions showed higher variability in controls (**Supplementary Fig. 4**).

**Figure 2:**
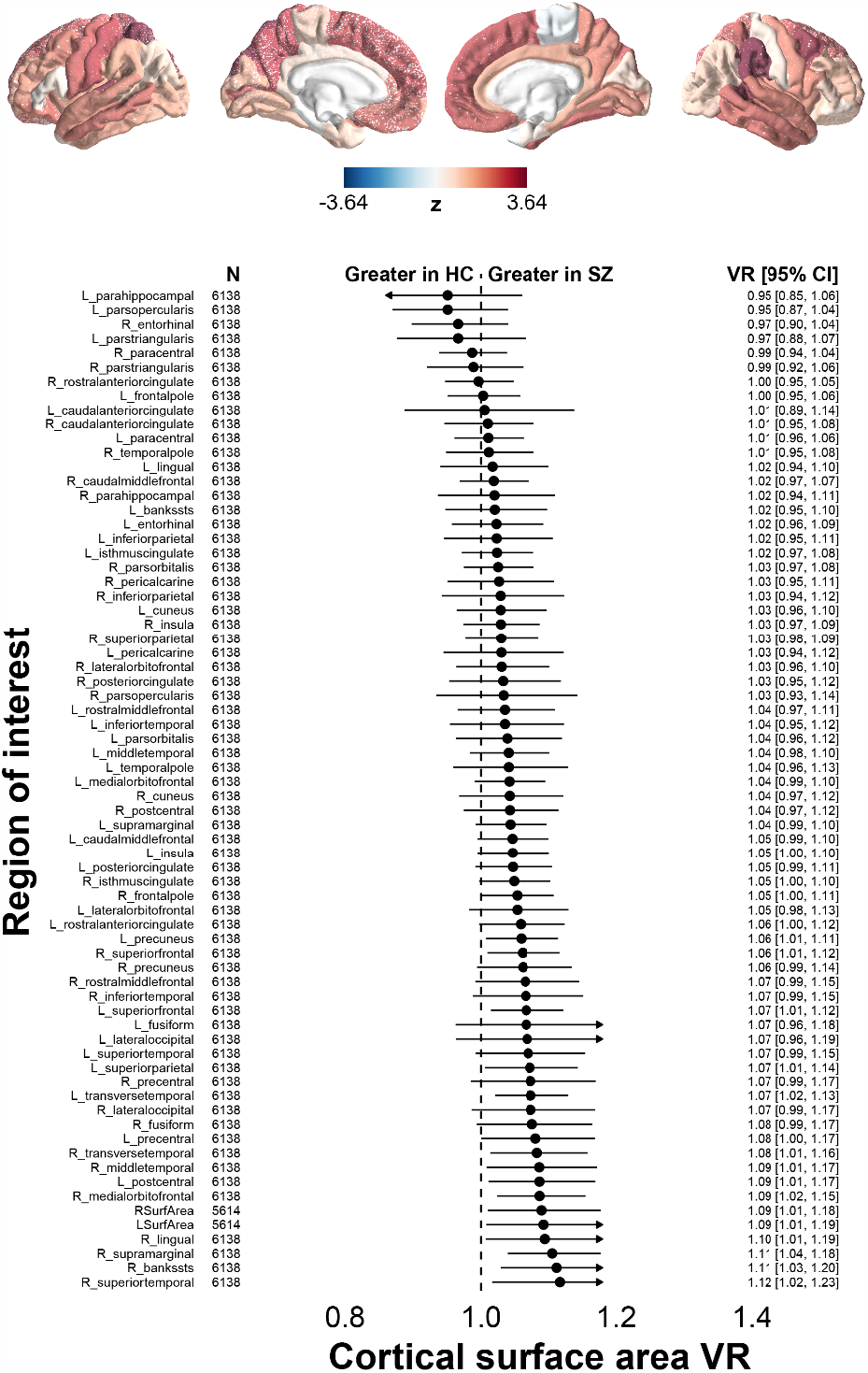
Variability ratio for cortical surface area. Same conventions are used as for Fig. 1. Higher heterogeneity in schizophrenia was observed for the following regions, irrespective of the deployed variability measure: Bilateral superiorfrontal and transverse temporal regions, cortical areas around the right superior temporal sulcus, right superiortemporal region, right supramarginal region, left postcentral region, right medialorbitofrontal region, left superiorparietal region, left precuneus, right middle temporal region and right lingual region.

### Cortical folding index

The schizophrenia group exhibited higher heterogeneity than controls in the right inferiortemporal region (VR = 1.33, 95% CI: 1.10 - 1.60) and lower heterogeneity in the left parahippocampal region (VR = 0.72, 95% CI: 0.56 - 0.91, **Fig. 3**). Higher heterogeneity of the right inferiortemporal region as well as higher homogeneity in the left parahippocampal region in schizophrenia were preserved when the coefficient of variation ratio was examined instead of the variability ratio (**Supplementary Fig. 5**). Of all gray matter measures, only the cortical folding index showed higher regional homogeneity in addition to higher regional heterogeneity in individuals with schizophrenia compared to healthy controls.

**Figure 3:**
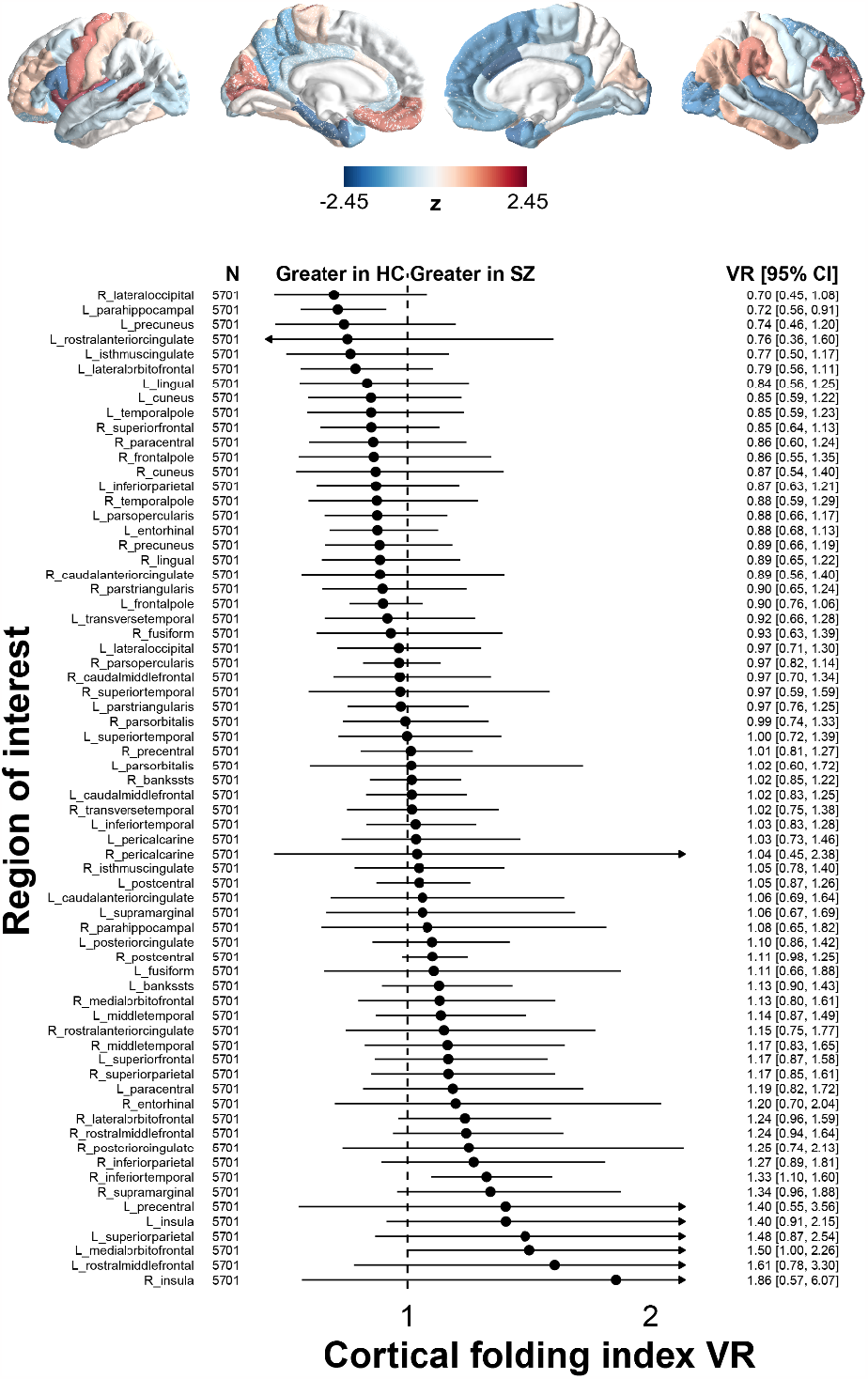
Variability ratio for cortical folding index. Same conventions as for Fig. 1. Irrespective of the applied variability measure, schizophrenia patients exhibited higher heterogeneity in the right inferiortemporal region, but lower heterogeneity in the left parahippocampal area.

### Subcortical volume

Bilaterally, the variability of the lateral ventricle volume and the volume of the inferior lateral ventricle was distinctly higher in the schizophrenia group than in the control group. Among the ventricle measurements, the highest effect size was found in the left inferior lateral ventricle (VR = 1.43, 95% CI: 1.26 - 1.61, **Fig. 4**). Additionally, higher variability in the bilateral pallidum and putamen, and in the left nucleus accumbens, left caudate and right hippocampus were evident in the schizophrenia compared to the control group. Among these brain regions, the highest effect sizes were found for the left nucleus accumbens (VR = 1.13, 95% CI: 1.04 - 1.22) and the left pallidum (VR = 1.10, 95% CI: 1.04 - 1.17, **Fig. 4**). No subcortical region showed significantly higher heterogeneity in the control group (**Fig. 4**). When the coefficient of variation ratio was computed instead of the variability ratio, higher heterogeneity in schizophrenia was confirmed for all regions except for bilateral pallidum and putamen (**Supplementary Fig. 6**). As for the variability ratio, no subcortical region exhibited higher heterogeneity in controls when the coefficient of variation ratio was examined (**Supplementary Fig. 6**).

**Figure 4:**
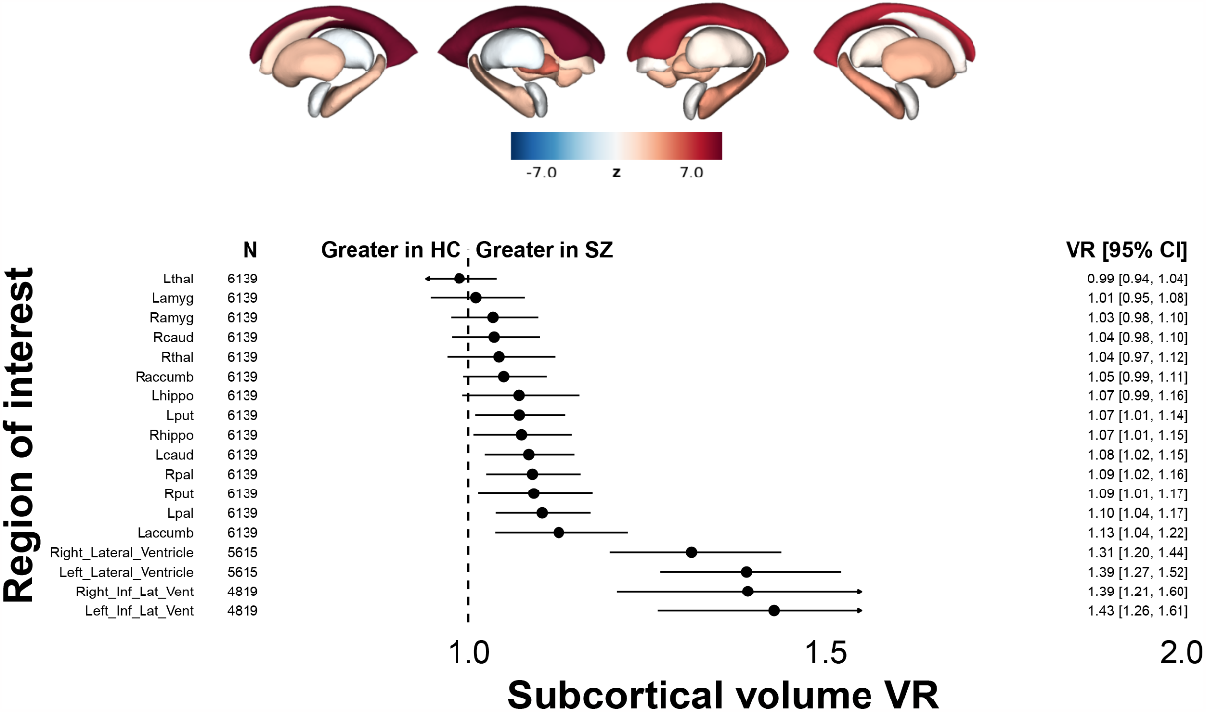
Variability ratio for subcortical volume. Same conventions as for Fig. 1. Lower panel: Variability ratio (VR) effect sizes are shown for subcortical volumes. Upper panel: VR effect sizes are projected onto respective subcortical structures and color-coded as z-values. In these subcortical maps, areas with higher VRs in patients appear in gradients of red, and areas with higher VRs in controls appear in gradients of blue. Independent of the applied variability measure, schizophrenia patients showed higher heterogeneity for lateral ventricles, left nucleus accumbens, left caudate and right hippocampus.

### Fractional anisotropy

Variability of the superior fronto-occipital fasciculus was lower in the schizophrenia compared to the control group, on both the left (VR = 0.86, 95% CI: 0.78 - 0.94) and right side (VR = 0.85, 95% CI: 0.75 - 0.95, **Fig. 5**). In contrast, no white matter tract showed greater heterogeneity in variability ratio in individuals with schizophrenia compared to controls. When the coefficient of variation ratio was examined instead of the variability ratio lower bilateral heterogeneity of the superior fronto-occipital fasciculus in the schizophrenia compared to the control group remained significant (**Supplementary Fig. 7**), and several regions including the left uncinate fasciculus exhibited greater heterogeneity in the schizophrenia compared to the control group (CVR = 1.16, 95% CI: 1.01 - 1.33, **Supplementary Fig. 7**).

**Figure 5:**
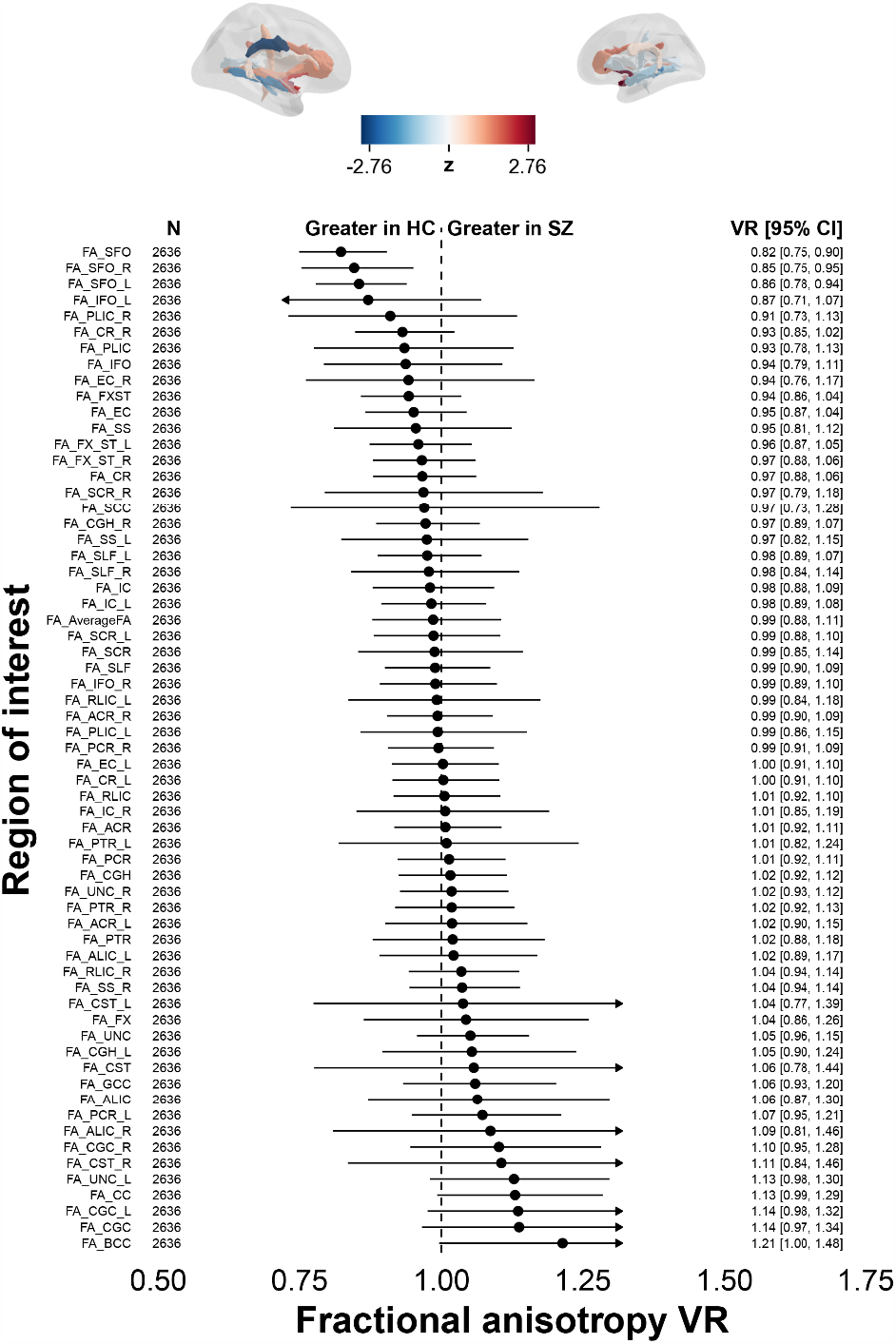
Variability ratio for fractional anisotropy. Same conventions as for Fig. 1. *Lower panel*: Variability ratio (VR) effect sizes are shown for fractional anisotropy (FA) of white matter structures. *Upper panel*: VR effect sizes are projected onto respective white matter structures and color-coded as z-values. In these maps, areas with higher VRs in patients appear in gradients of red, and areas with higher VRs in controls appear in gradients of blue. Irrespective of the deployed variability measure, higher homogeneity in schizophrenia was observed for the superior fronto-occipital fasciculus in both hemispheres.

### Person-based similarity index

At a global level, the person-based similarity index (PBSI) analysis revealed that heterogeneity of cortical thickness is similar for schizophrenia patients and healthy controls (p = 4.891e-01, Mann-Whitney U test, Bonferroni-corrected, **Supplementary Fig. 8**). However, schizophrenia patients showed less heterogeneity (higher PBSI) in cortical surface area, subcortical volume and especially in the folding index compared to healthy controls (folding index: p = 2.123e-16; surface area: p = 4.972e-06; subcortical volume: p = 8.33e-06, Mann-Whitney U test, Bonferroni-corrected, **Supplementary Fig. 8**). On the other hand, individuals with schizophrenia exhibited higher heterogeneity (lower PBSI) in fractional anisotropy (p = 2.649e-09, Mann-Whitney U test, Bonferroni-corrected, **Supplementary Fig. 8**).

## Discussion

The results of this comprehensive meta- and mega-analysis extend our understanding of structural brain heterogeneity in schizophrenia, an aspect that has been insufficiently studied to date, yet holds significant implications for the identification of illness subtypes and informative biomarkers^7^. A strength of this study was the comprehensive heterogeneity assessment of multiple structural brain measures based on an ENIGMA Schizophrenia Working Group dataset, a large international repository of MRI-based brain measures collected from various sites globally^11,12,23^. Different structural brain measures are thought to be influenced by separate sets of genes during different stages of development^34–39^, and the multimodal assessment and comparison of structural brain measures is instrumental to disentangle their potentially different alterations in schizophrenia^13^. This study therefore investigated and compared the variability in cortical thickness, surface area, folding index, subcortical volume, and fractional anisotropy, providing a thorough assessment of regional and global variability. Moreover, the framework of the large ENIGMA dataset increases the robustness of the results and therefore the replicability of findings in psychiatry and neuroscience^23,40–44^.

One of the key findings at the regional level is greater multimodal variability in schizophrenia for frontotemporal structures. While cortical surface area and folding index were more heterogeneous in a few temporal and frontal regions, higher variability of cortical thickness encompassed widespread bilateral temporal areas in addition to some frontal regions. Thus, these results suggest that schizophrenia is associated with multimodal structural variability in different components of a frontotemporal network, thereby extending prior findings that show on average thinning of frontal and temporal regions in schizophrenia^12,13,45,46^. A possibility would be that different subgroups exist that can be distinguished by different neuroanatomical correlates: While one subgroup may have structural irregularities in temporal rather than frontal regions, be it cortical thinning, folding abnormalities or a combination thereof, another subgroup may be characterized by the inverse structural pattern. In all cases, the result would likely be impaired frontotemporal interaction, a neurobiological hallmark of psychotic disorders. Further investigation of neuroanatomical features with regard to potential subgroups could therefore be an interesting route for future studies. At least with regard to mean-scaled variability, schizophrenia patients also showed greater heterogeneity of fractional anisotropy in the left uncinate fasciculus which connects temporal with frontal regions. Structural deficits of the left uncinate fasciculus may therefore also contribute to aberrant frontotemporal interaction in a subset of patients. On the other hand, our study also showed that gyrification of the left parahippocampal region is more homogeneous in schizophrenia compared to controls. Gyrification characteristics of the left parahippocampal region may therefore occur uniformly across potential subtypes and represent a central attribute of the disorder. Interestingly, a part of the left parahippocampal region has been recently shown to be thinner and associated with hallucinations in schizophrenia spectrum individuals^47^. In our study, greater variability was also observed with regard to the volume of the left nucleus accumbens and caudate, regions implicated in reward processing and goal-directed motor control, respectively^48,49^. Variability in these subcortical structures may be linked to the heterogeneity of negative and catatonic symptoms that differ substantially across schizophrenia individuals^50,51^. Overall, the largest effect sizes with regard to greater variability ratio were observed for the lateral ventricles in both hemispheres. Since ventricular enlargement is among the best-established neurobiological findings in schizophrenia patients^7,11,52–55^, it is notable that higher variability in schizophrenia remained robust when the coefficient of variation - which accounts for group differences in mean volumes - was used instead of the variability ratio. In contrast, a previous study comparing first-episode schizophrenia with healthy controls showed higher variability of lateral ventricles only for the variability ratio, but not for the coefficient of variation^7^. This difference with our study could be related to the longer illness duration in our study, which included individuals with chronic schizophrenia.

Notably, at a global level, a part of our findings challenges the prevailing view of greater brain heterogeneity in schizophrenia in comparison to healthy individuals^7,14,15,18^. The person-based similarity index (PBSI) analysis^16,17^ surprisingly revealed that individuals with schizophrenia are characterized by a significantly higher homogeneity of especially the folding index, a measure related to the gyrification of the brain. This observation supports the view that certain gyrification characteristics could represent a potential core feature of schizophrenia^39^. Previous studies demonstrated that adolescent and young adult individuals with schizophrenia typically feature stronger gyrification in widespread regions compared with healthy controls^56–66^, possibly related to overexpression of fibroblast growth factor-2 gene in the dorsolateral prefrontal cortex and further molecular-level aberrations in schizophrenia^39,67^. While gyrification patterns are determined early in development^68^, it is plausible that certain folding characteristics might render the brain more susceptible to the later development of schizophrenia, thus presenting a potential predictive biomarker for the disorder. In line with this view, previous studies with limited sample size have shown that schizophrenia is associated with aberrant gyrification characteristics in specific cortical regions which are in part related to particular symptoms and deficits of the disorder^58,59,62,63,65,69–73^. For instance, the severity of positive symptoms in schizophrenia has been associated with gyrification in the right frontal and temporolimbic region^63,64^. Similarly, fronto-temporo-limbic disconnectivity in schizophrenia, which is related to delusions, hallucinations and disorganization symptoms^74,75^, has been proposed to be linked with local gyrification patterns^39^. Previous studies also compared PBSI scores for cortical thickness, subcortical volume and cortical surface area between schizophrenia or first episode psychosis patients and healthy controls^15,16^. While some results are in line with ours, e.g. higher cortical surface area PBSI scores in psychosis patients^16^, others differ, e.g. higher cortical thickness PBSI scores in controls^15,16^ instead of no significant difference in our study. Deviations from our results may be related to the more limited sample size in both studies as well as the shorter illness duration of first-episode psychosis patients in Antoniades et al^16^. However, the PBSI analysis also revealed that individuals with schizophrenia exhibit greater global heterogeneity than controls with regard to fractional anisotropy, suggesting that different white matter abnormalities contribute to schizophrenia subtypes. White matter pathology has been associated with functional dysconnectivity in schizophrenia^76^, and resulting impairment of neural network synchrony may underlie schizophrenia symptoms^77,78^.

Despite the findings, this study is not without limitations. Not all data were individual-level data, which could potentially introduce bias. Moreover, not all measures were collected at all sites, and in particular the amount of fractional anisotropy data was lower in comparison to other measures. In addition, the parcellation of gray and white matter networks was coarse, and a more fine-grained parcellation in future studies may lead to more differentiated insights of structural variability.

In conclusion, this study provides novel insights into the heterogeneity of structural brain measures in schizophrenia. In light of increasing interest in schizophrenia subtypes and corresponding neurobiological characteristics^79–91^, high variability in certain neuroanatomical structures may point to potential avenues for the development of more targeted treatment strategies. On the other hand, low variability of structural features such as gyrification may substantiate core features of schizophrenia that are shared across illness subtypes. Our results hold potential to guide future research efforts and pave the way for precision medicine approaches in schizophrenia. Ultimately, this study underscores the critical role of variability in our understanding of complex disorders like schizophrenia, a perspective that is essential for a more holistic approach to its neurobiology.

## Acknowledgements

We thank Dr. Ellen Ji for her scientific advice and her support in data management.

## Author contributions

W. Omlor co-analyzed and interpreted the data, co-performed the statistical analysis, wrote the first draft of the manuscript and revised the manuscript. S. Fuchs and F. Rabe co-analyzed the data, and S. Fuchs additionally took care of data reception and management. P. Homan initiated, conceptualized and supervised the study, co-analyzed the data and co-performed the statistical analysis. All other authors provided a part of the dataset and refined the manuscript. All authors approved the final version of the manuscript.

## Supplementary Information

**Supplementary figure 1:**
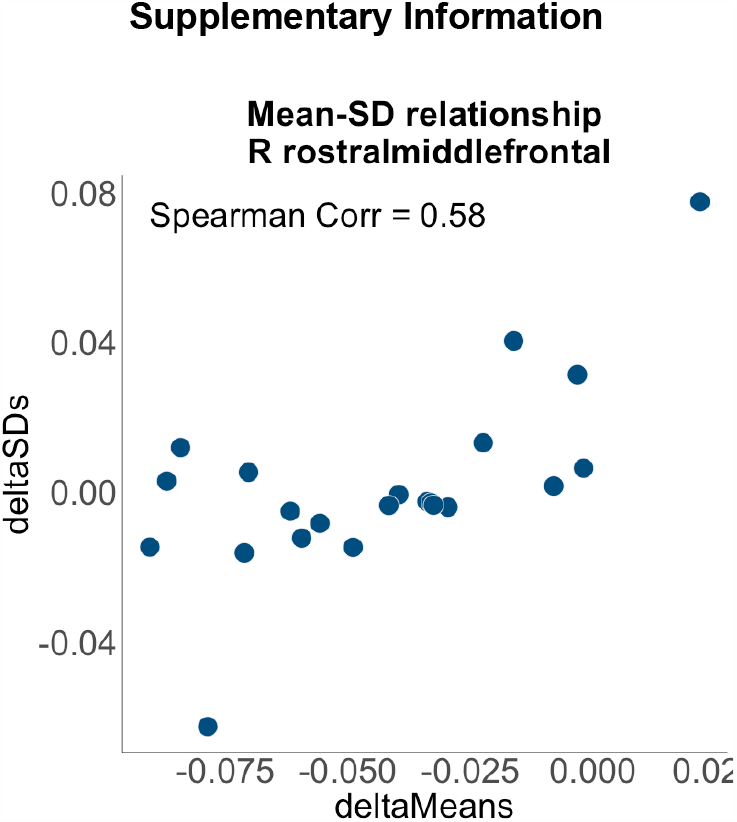
Relationship between mean and standard deviation across sites - cortical thickness of the right rostral middle frontal cortex (R rostralmiddlefrontal). Differences of standard deviations (deltaSDs) are plotted against differences of means (deltaMeans). Difference values were obtained by substracting Mean/SD of controls from Mean/SD of patients. For cortical thickness, the right rostral middle frontal cortex showed the highest positive correlation between deltaMeans and deltaSDs.

**Supplementary figure 2:**
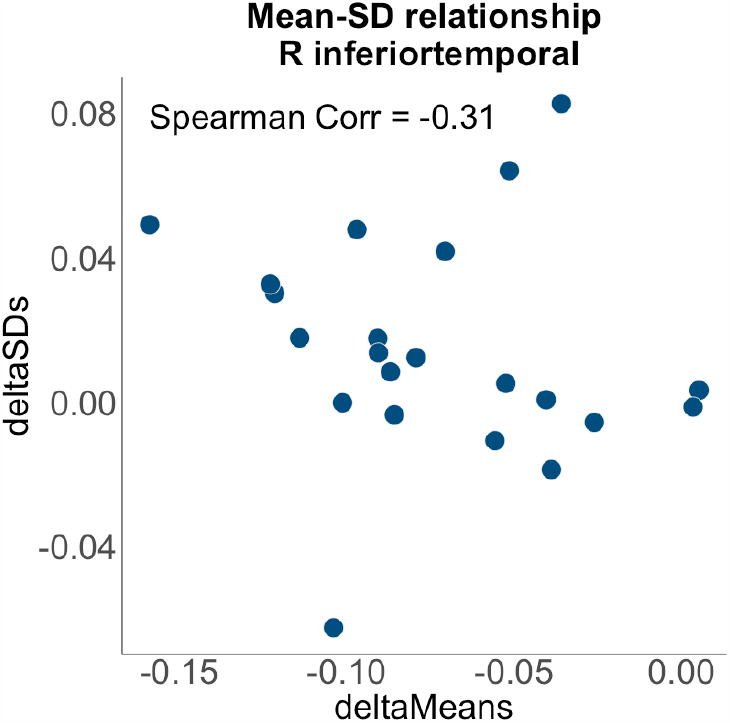
Relationship between mean and standard deviation across sites - cortical thickness of the right inferior temporal cortex (R inferiortemporal). Same conventions as for Supplementary Figure 1. For cortical thickness, the right inferior temporal cortex showed the highest negative correlation between deltaMeans and delta SDs.

**Supplementary figure 3:**
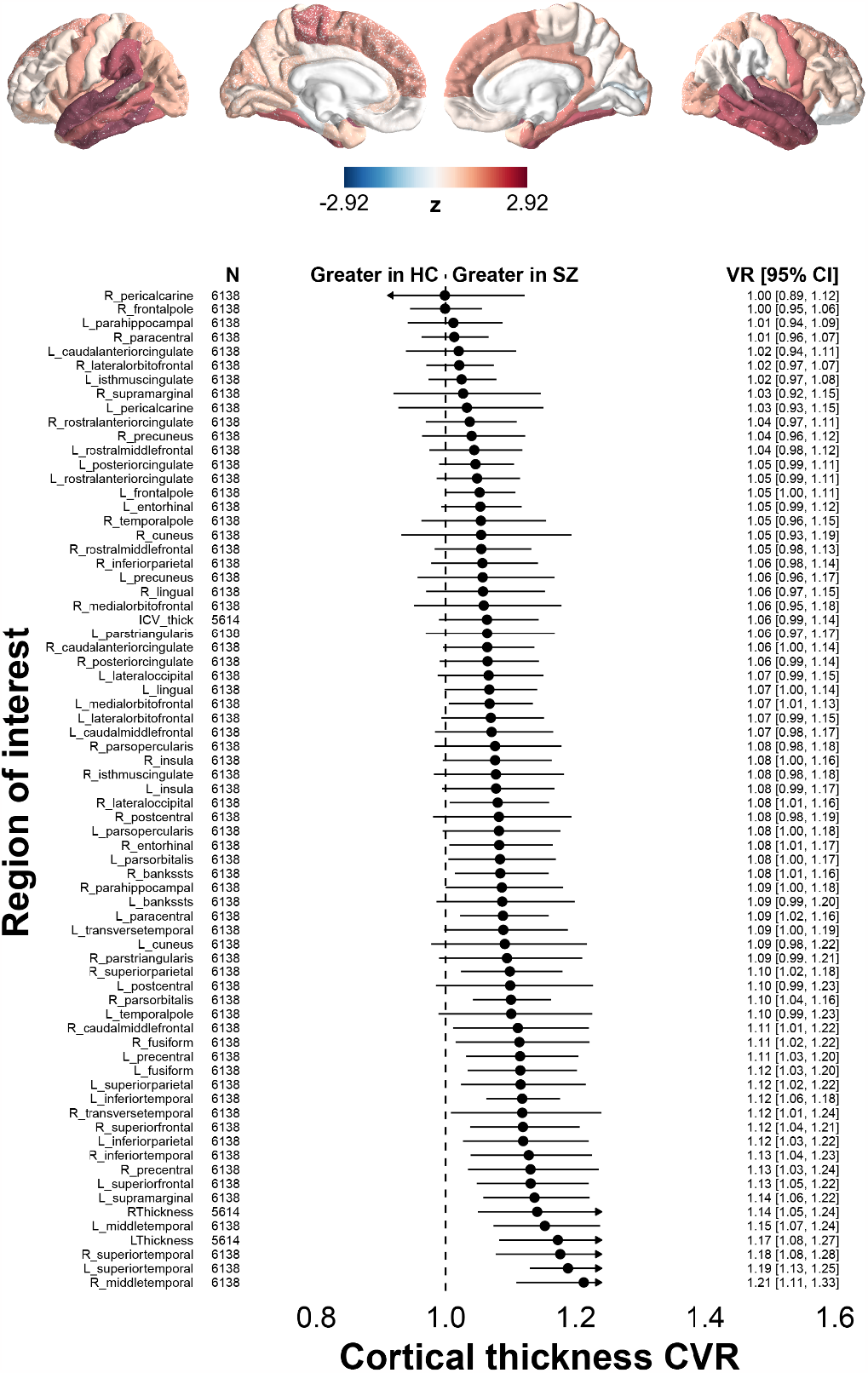
Coefficient of Variation ratio for cortical thickness. Lower panel: Coefficient of variation ratio (CVR) effect sizes for schizophrenia patients (SZ) vs. healthy controls (HC) are shown for different cortical regions and on a linear scale, statistically controlling for age and gender. Upper panel: CVR effect sizes are projected onto the brain surface and color-coded as z-values. In these cortical maps, areas with higher CVRs in patients appear in gradients of red, and areas with higher CVRs in controls appear in gradients of blue. CI: Confidence Interval.

**Supplementary figure 4:**
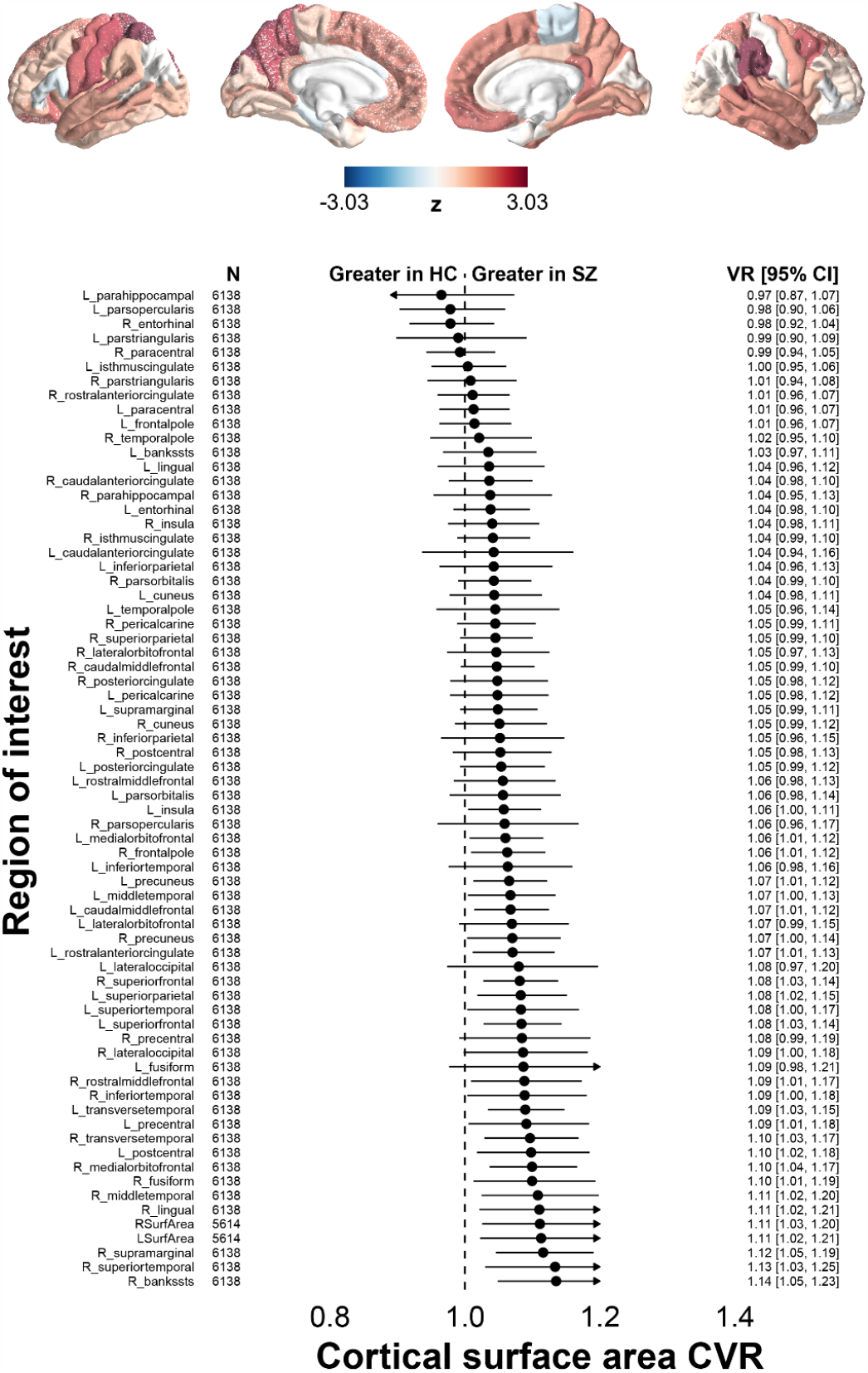
Coefficient of Variation ratio for cortical surface area. Same conventions as for Supplementary Fig. 3.

**Supplementary figure 5:**
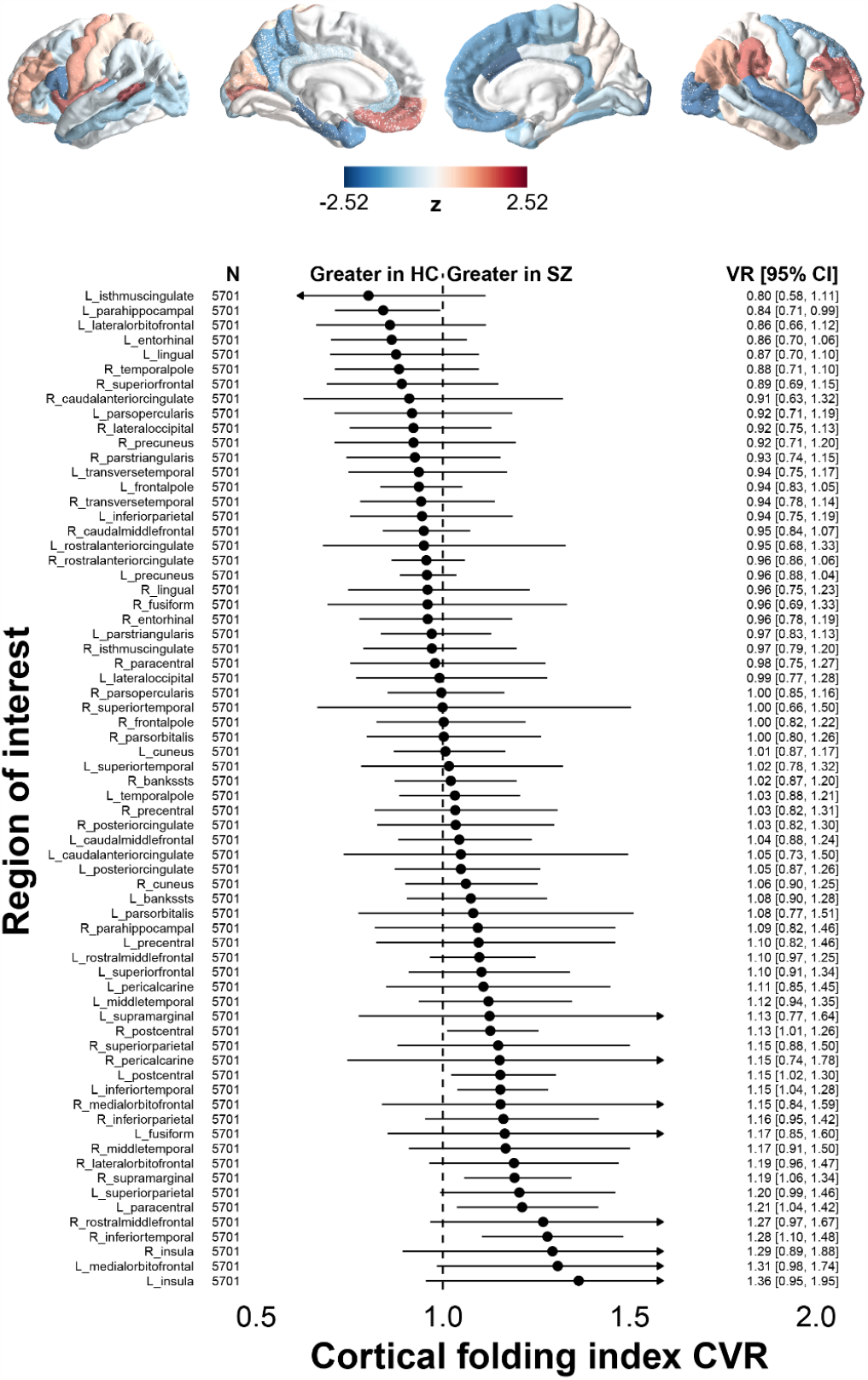
Coefficient of Variation ratio for cortical folding index. Same conventions as for supplementary Fig. 3.

**Supplementary figure 6:**
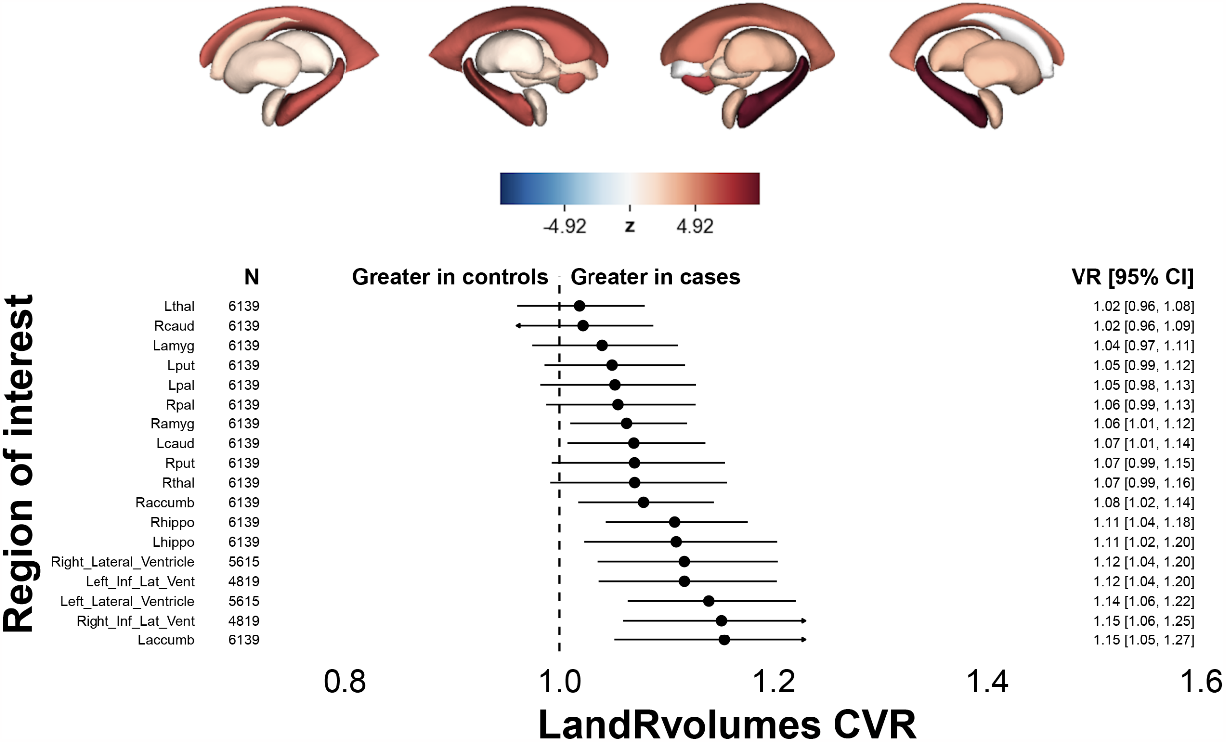
Coefficient of Variation ratio for subcortical volume. Same conventions as for supplementary Fig. 3. Lower panel: Coefficient of Variation ratio (CVR) effect sizes are shown for subcortical volumes on the left and right side (LandRvolumes). Upper panel: CVR effect sizes are projected onto respective subcortical structures and color-coded as z-values. In these subcortical maps, areas with higher CVRs in patients appear in gradients of red, and areas with higher CVRs in controls appear in gradients of blue.

**Supplementary figure 7:**
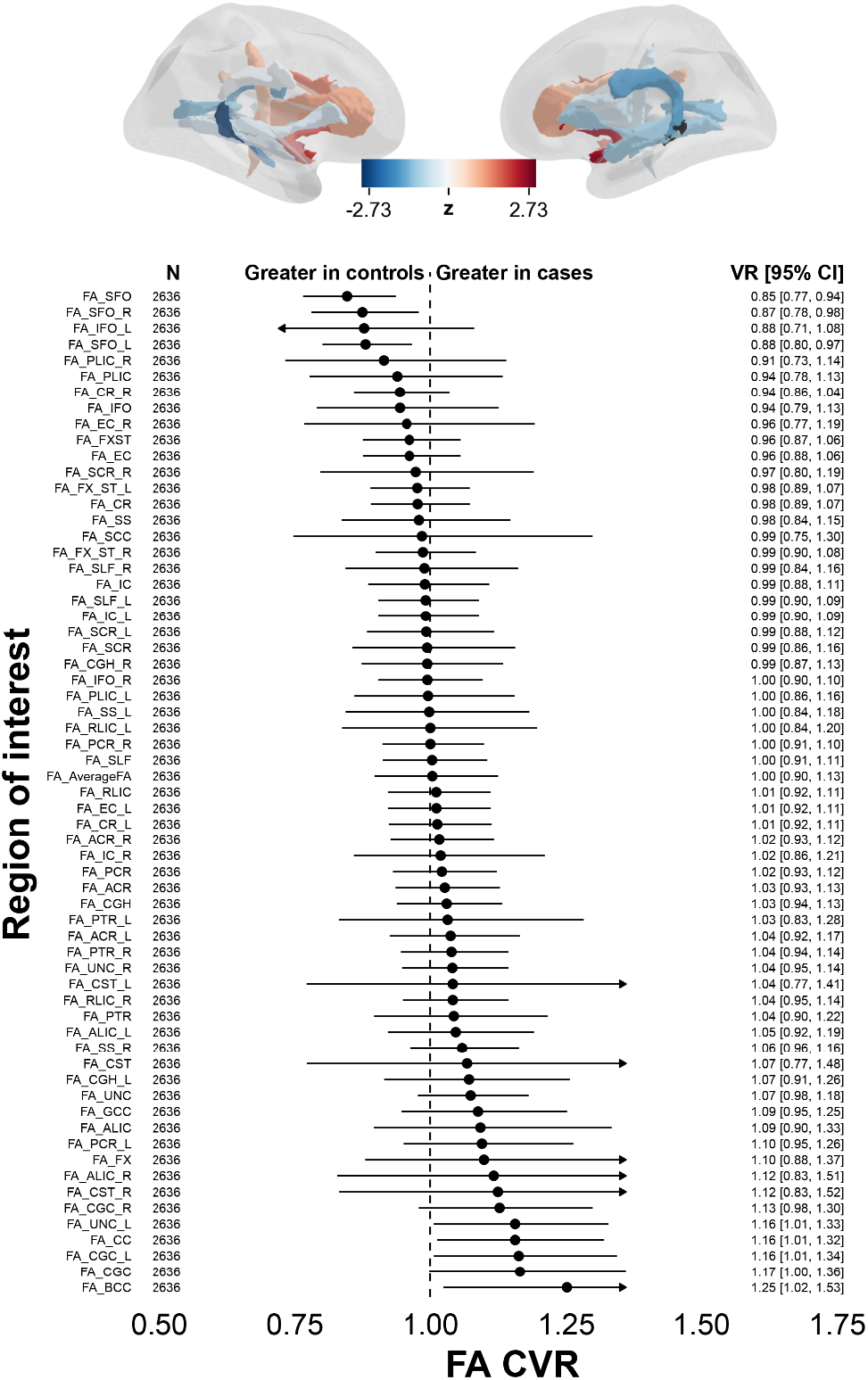
Coefficient of Variation ratio for fractional anisotropy. Same conventions as for supplementary Fig. 3. Lower panel: Coefficient of Variation ratio (CVR) effect sizes are shown for fractional anisotropy (FA) of white matter structures. Upper panel: CVR effect sizes are projected onto respective white matter structures and color-coded as z-values. In these maps, areas with higher CVRs in patients appear in gradients of red, and areas with higher CVRs in controls appear in gradients of blue.

**Supplementary figure 8:**
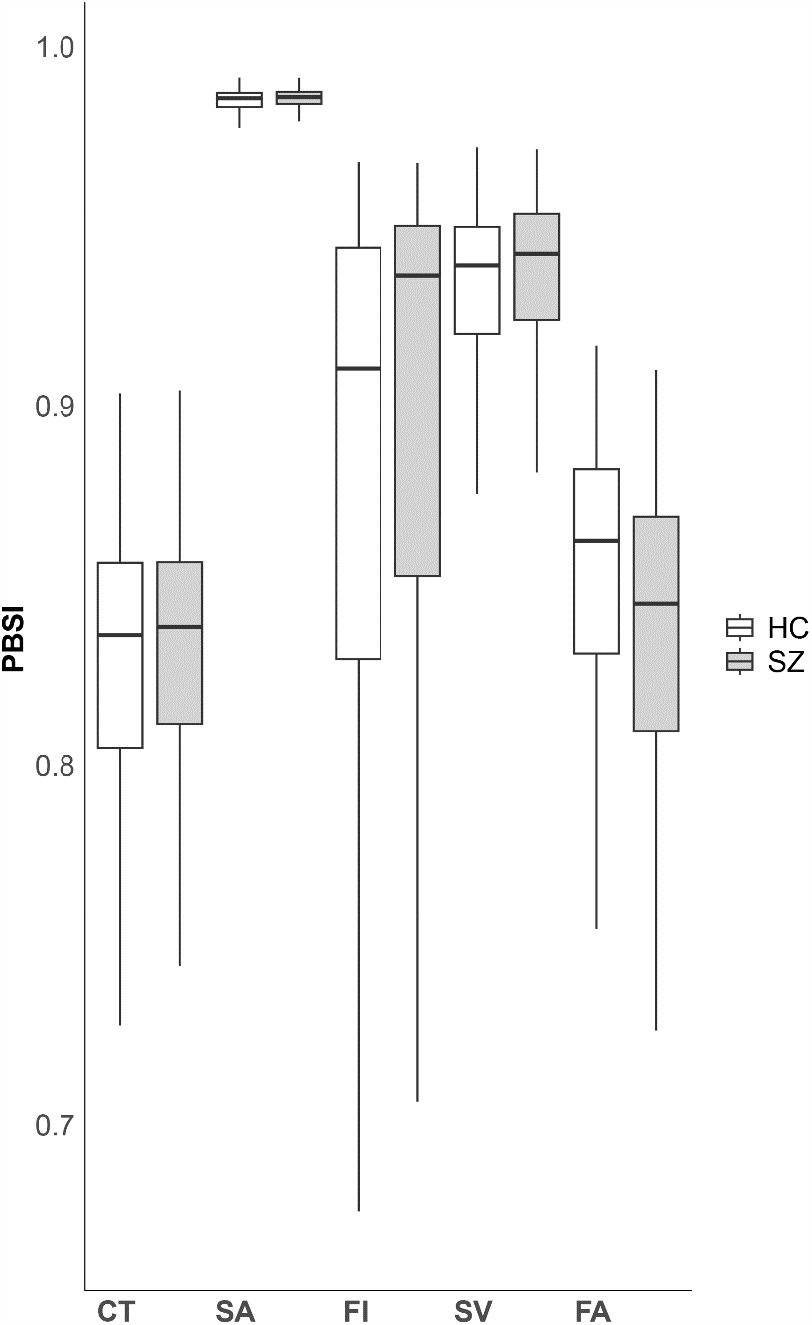
Person-based similarity index. Person-based similarity index (PBSI) for cortical thickness (CT), cortical surface area (SA), cortical folding Index (FI), subcortical volume (SV) and fractional anisotropy (FA).

**Supplementary Table 1:** Sample demographics.

| Site | Level | Patients (T1) | Controls (T1) | Patients (DTI) | Controls (DTI) |
| --- | --- | --- | --- | --- | --- |
| ASRB | Group | 263 | 166 | 326 | 197 |
| CASSI | Individual | 58 | 65 | 0 | 0 |
| CIAM | Individual | 19 | 31 | 12 | 25 |
| COBRE | Individual | 73 | 70 | 73 | 70 |
| ESO | Individual | 40 | 40 | 0 | 0 |
| FBIRN | Group | 184 | 174 | 145 | 144 |
| FIDMAG | Individual | 57 | 57 | 57 | 57 |
| HIULONG | Individual | 245 | 88 | 245 | 88 |
| IGP | Group | 68 | 71 | 0 | 0 |
| IMH | Individual | 178 | 111 | 0 | 0 |
| IRCCS | Individual | 172 | 116 | 78 | 144 |
| MPRC | Individual | 230 | 270 | 0 | 0 |
| OLIN | Individual | 135 | 389 | 0 | 0 |
| OSAKA1 | Group | 142 | 402 | 0 | 0 |
| OSAKA2 | Group | 76 | 237 | 63 | 229 |
| OSAKA3 | Group | 82 | 413 | 82 | 327 |
| PUK | Individual | 60 | 28 | 0 | 0 |
| SWIFT | Individual | 24 | 14 | 0 | 0 |
| TOP | Individual | 352 | 756 | 170 | 582 |
| UCISZ | Individual | 26 | 29 | 0 | 0 |
| UNINA | Individual | 49 | 55 | 49 | 55 |
| Paris | Group | 33 | 55 | 0 | 0 |

